# Hybrid Refinement of Heterogeneous Conformational Ensembles using Spectroscopic Data

**DOI:** 10.1101/656595

**Authors:** Jennifer M. Hays, David S. Cafiso, Peter M. Kasson

## Abstract

Multi-structured biomolecular systems play crucial roles in a wide variety of cellular processes but have resisted traditional methods of structure determination which often resolve only a few low-energy states. High-resolution structure determination using experimental methods that yield distributional data remains extremely difficult, especially when the underlying conformational ensembles are quite heterogeneous. We have therefore developed a method to integrate sparse, multi-multimodal spectroscopic data to obtain high-resolution estimates of conformational ensembles. We have tested our method by incorporating double electron-electron resonance data on the SNARE protein syntaxin-1a into biased molecular dynamics simulations. We find that our method substantially outperforms existing state-of-the-art methods in capturing syntaxin’s open/closed conformational equilibrium and further yields new conformational states that are both consistent with experimental data and may help in understanding syntaxin’s function. Our improved methods for refining heterogeneous conformational ensembles from spectroscopic data will greatly accelerate the structural understanding of such systems.

---

Heterogeneous conformational ensembles play important roles in in a wide variety of cellular processes and diseases, including flexible recognition events during infection and in signal transduction pathways.^1–6^ The major challenges in structural refinement of these flexible systems are twofold. Often, experimental data yield ensemble-average quantities and/or the experimental data are sparse.^7–11^ Both of these difficulties lead to an ill-posed inverse problem: in both cases, an ensemble of structures, which is degenerate in the experimental quantity of interest, must be enumerated with very little other information.^12–14^ One way to avoid this inverse problem is to use a forward model, such as an MD forcefield, and directly integrate the experimental data into the model such that the final ensemble reproduces the experimental data. A great deal of work has been done to develop methods for integrating ensemble-average quantities into forward models and this work has been quite successful.^15–22^ However, a robust strategy for integrating sparse, distributional data has remained elusive.

Here we describe a hybrid maximum-entropy/stochastic-resampling approach for biasing molecular simulation ensembles towards experimental distributions rather than ensemble averages. We apply this method to double electron-electron resonance (DEER) data, but the method is extremely general and may be used for nearly any experimental method yielding distributional data. The method exhibits no instabilities in regions of zero probability and can sample important backbone conformational change. We describe how to incorporate a single distribution first, then discuss a simple generalization to multiple distributions.

Our hybrid approach, which we call bias-resampling ensemble refinement (BRER), uses an iterative refinement scheme to update an estimate of the conformational ensemble 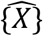 based on a DEER distribution *P*_DEER_(*d*) (Fig. 1). The simplest formulation of BRER is described here, while a more complex variant is given in the Supplement. During refinement, each conformation 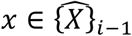 is updated using a biased MD simulation such that the updated estimate 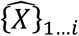 better reproduces *P*_DEER_(*d*). Thus, over the course of multiple rounds of refinement, the conformational estimate 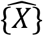 should yield a distribution 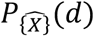 that converges on *P*_DEER_(*d*). The initial estimate 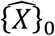 may be obtained using experimental data (an NMR ensemble, a single crystal structure) or an experimentally-informed model.

**Figure 1.**
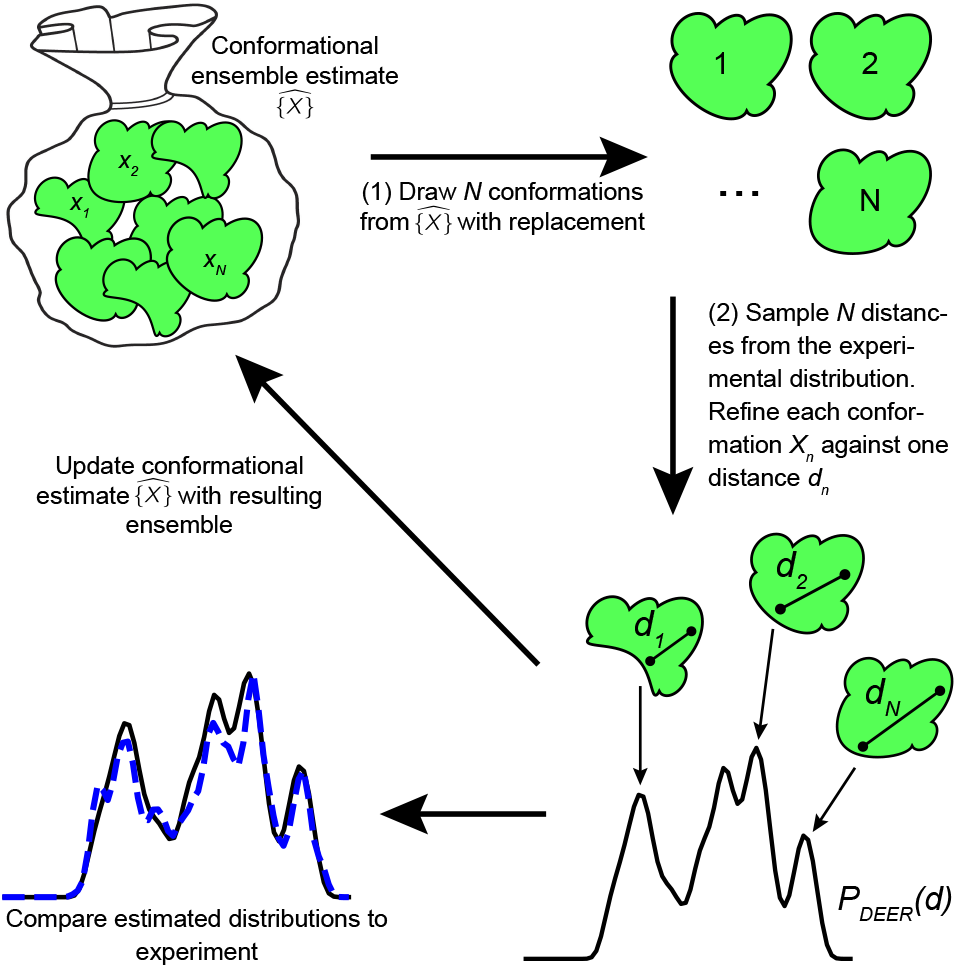
Estimation of a conformational ensemble {*X*} by stochastic resampling of experimental data. An iterative update framework for bias resampling ensemble refinement from an initial estimate is schematized as follows: (1) a set of *N* conformations is drawn from the conformational ensemble estimate 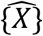 and (2) each conformation is refined against a single target which is stochastically resampled from an experimental distribution. At each iteration, the estimated distribution calculated from 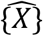 is compared against the experimental distribution. If the distribution 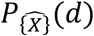 is significantly different from *P*_experimental_(*d*), the conformational estimate 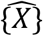 is updated with the refined structures and refinement procedure is repeated.

The stochastic-resampling approach is implemented as follows. Because any distribution *P*_DEER_(*d*) may be represented as a sum of Gaussians, let the experimental distribution be a linear combination of *M* Gaussians with centers located at *d*_*m*_:

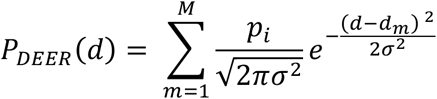

with weighting factor *p*_*i*_ During the *i*^th^ round of refinement, we randomly sample a set of *N* structures from the conformational estimate of the previous round, 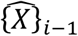. Each structure is assigned one target distance *d*_*n*_ via a probability-weighted draw from the set of experimental distances *{d_m_}*. A maximum-entropy biasing potential is then applied to each ensemble member, driving the member towards its target distance.^23–25^ We allow the ensemble to relax at the target distribution (which has been resampled from *P*_DEER_(*d*)) for some time *t*, at which point the resampling procedure is repeated. Because this approach is equivalent to performing Monte Carlo with an acceptance probability of one, the simulation ensemble distribution should converge on the experimental distribution after sufficient repetitions of the resampling procedure.

The two components of this method are demonstrated separately in Figure 2. In Fig 2A, we show how stochastic resampling converges on a complex target distribution demonstrated using a Gaussian stub in place of the biased MD. Figure 2B shows the results of an MD simulation biased to a single target. The biasing potential successfully drives the simulation distance to the target distance without disrupting secondary structural elements experimentally known to be preserved.^26^ Details of the biased MD are provided in the Supplement.

**Figure 2.**
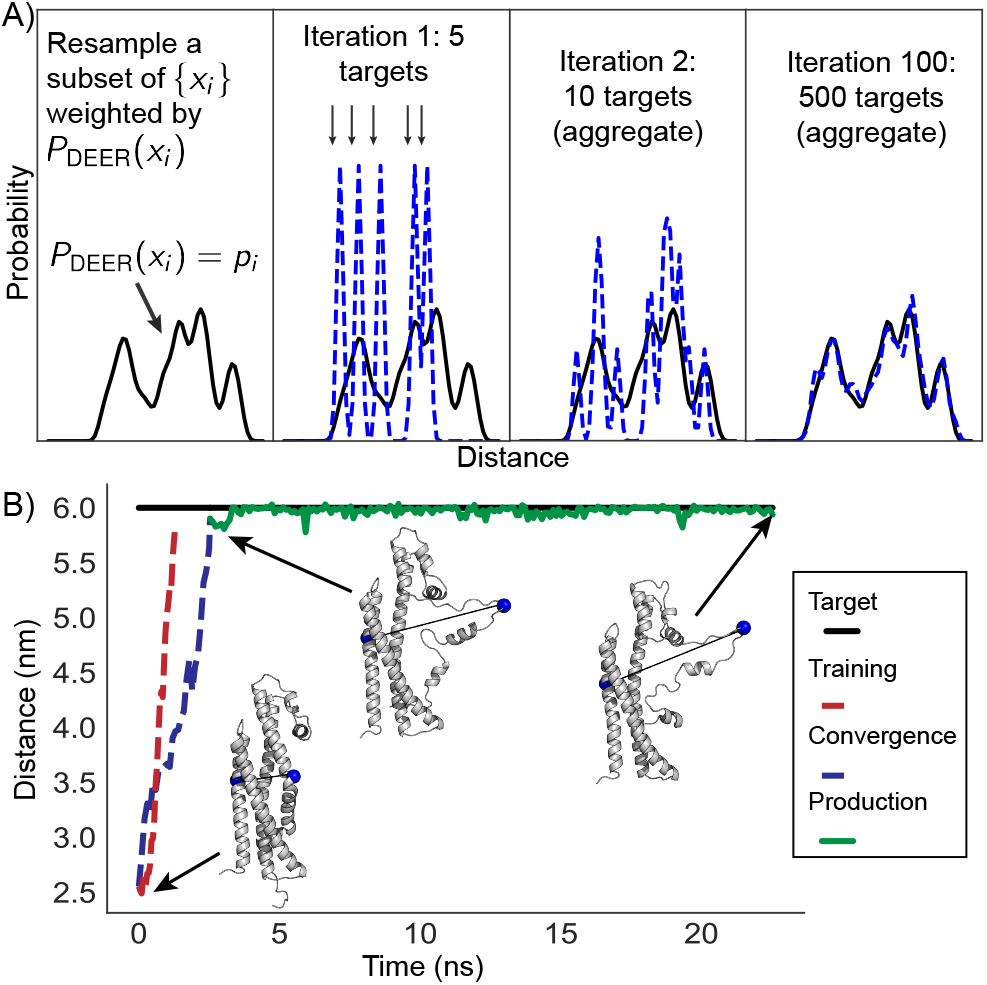
Bias-resampling ensemble refinement has two components: stochastic resampling and a maximum-entropy biasing potential. Iterative stochastic resampling of the target distribution yields an excellent approximation after 500 targets have been drawn (A). Here, the more complex MD engine has been replaced with a Gaussian stub such that each ensemble member samples a simple Gaussian distribution around its target. An example of the maximum-entropy biasing potential for a single target is shown in (B). First, a maximum-entropy coupling constant is trained, which ensures that a minimally perturbative quantity of energy is introduced into the system. The simulation is then restarted using the pre-trained coupling constant and converges to its target without excessively disrupting the secondary structure of the biased conformation.

This method is trivially extensible to multiple distributions since resampling can be performed on a joint distribution. If information on the correlation structure of the distributions is unavailable, they are assumed to be independent, and draws are performed on the convolution of the distributions. Our approach is especially powerful since information about correlation structure can be recovered from the ensemble; because the coupling constants are first trained using a maximum-entropy formalism, we can measure the work needed to drive the ensemble to its target distances. This quantity reports on the correlation between the particular distributional modes that have been sampled. However, in this letter we focus on the effectiveness of the fundamental method.

We have used the BRER methodology to refine the conformational ensemble of the soluble domain of syntaxin1-a using previously published DEER data.^26^ We find that BRER substantially outperforms current state-of-the-art methods at reproducing the experimental distributions and identifies previously unknown structural sub-states. These sub-states suggest that the open state of syntaxin may be more conformationally diverse than previously thought and thus have direct implications for formation of the SNARE complex.

We obtained three experimentally-derived distributions from residue-residue pairs 52/210, 105/216, and 196/228 and used these distributions to refine the conformational ensemble of soluble syntaxin via three different methods: BRER, EBMetaD,^27^ and restrained-ensemble MD.^28^ The BRER-derived distributions reproduce the experimental distributions significantly better than either EBMetaD or restrained-ensemble (Fig. 3A), quantified by Jensen-Shannon divergence (Fig. 3B). BRER performs particularly well at reproducing the 52/210 distribution; the well-separated bimodal peaks in this distribution are important because they directly report on syntaxin’s open/closed equilibrium.^26,29–31^ Because EBMetaD and restrained-ensemble MD suffer from numerical instabilities in regions of zero probability, these methods fail to accurately reproduce this distribution. EBMetaD samples the open and closed states but with incorrect relative probabilities, and restrained-ensemble simulations simply fail to sample the open state. The 105/216 and 196/228 distributions pose a less challenging problem for EBMetaD and restrained-ensemble MD since neither distribution has very well-separated modes, yet they are still better reproduced by BRER. Thus, BRER is a top-performing, general method for refining conformational ensembles using DEER data.

**Figure 3.**
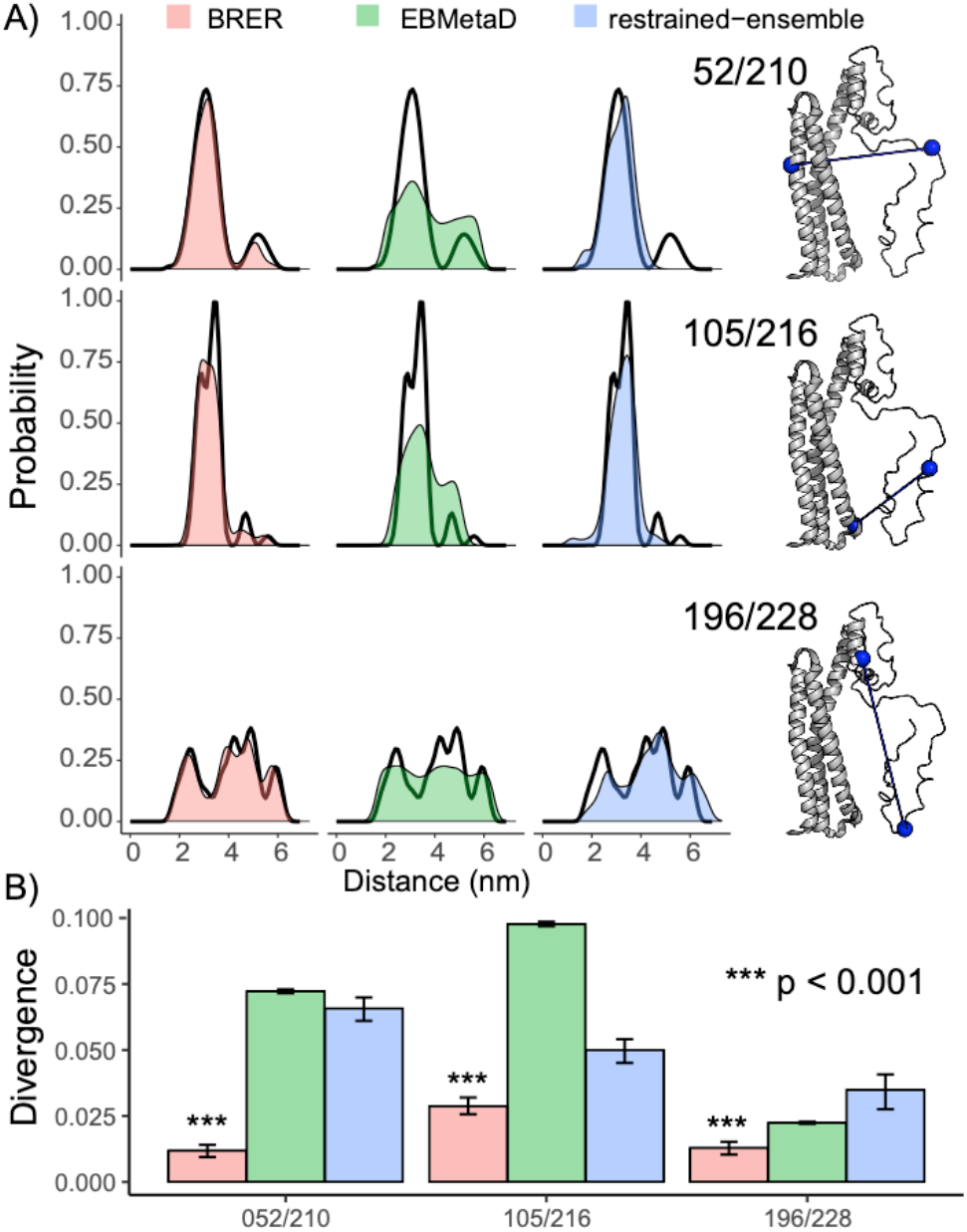
BRER outperforms current state-of-the-art methods for refining conformational ensembles against DEER distributions. Three sets of ensembles were refined against the three experimental distributions (shown in black in A). Distributions calculated from the BRER, EBMetaD, and restrained-ensemble conformational estimates are shown in color. BRER both qualitatively (A) and quantitatively (B) outperforms these other state-of-the-art methods for all three distributions. Agreement with the experimental distributions is quantified as Jensen-Shannon divergence (B).

Refinement of the syntaxin conformational ensemble with these pairs yields a previously unobserved family of structures that are partially open. It is generally thought that syntaxin must be in an open state to participate in formation of the SNARE complex and thus perform its critical role in neuronal exocytosis.^30–34^ No experimental structures of the apo syntaxin open state exist, but it has been hypothesized that the open state is characterized by complete dissociation of the H3 domain from the Habc domain and an unwinding of the linker region between Hc and H3 (Fig. 4A).^35–37^ The BRER-refined ensemble identifies additional structures in which the H3 domain is only partially dissociated from Habc and the linker region retains is secondary structure (Fig. 4B), with tight contacts remaining between residues 146-156 and 187-198. These BRER-refined structures are in close agreement with the DEER-derived distributions, as are structures in which H3 completely dissociates (Fig 4C). These results suggest a testable hypothesis: the syntaxin conformational ensemble is more diverse than was previously thought and both the partially open and fully open states contribute to the ensemble. In this scenario, formation of the SNARE complex could result from a further conformational selection process. Additional DEER experiments informed by the BRER-refined ensemble could elucidate whether the open-state ensemble is indeed conformationally diverse and whether a conformational selection process takes place to form the final SNARE bundle.

**Figure 4:**
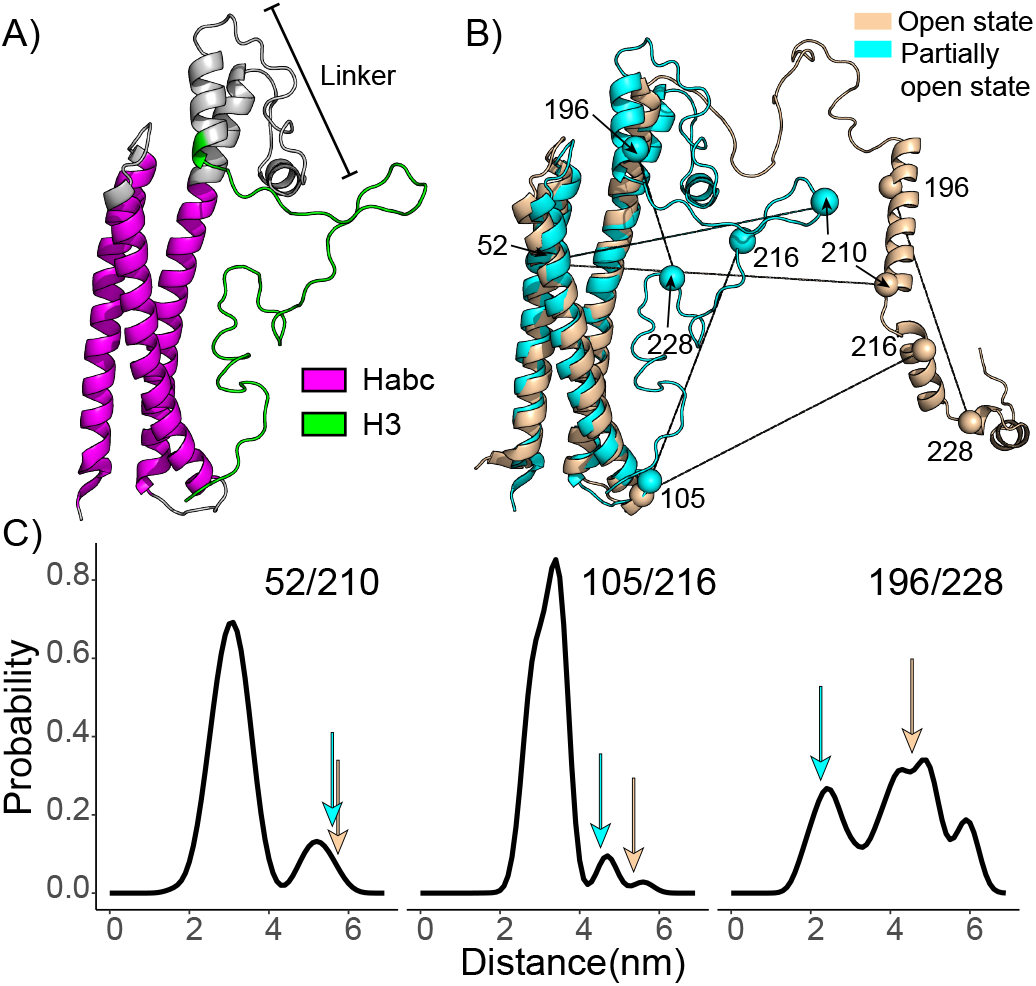
BRER-guided refinement yields previously unresolved conformational sub-states. Rendered in (A) is a new partially open conformation of syntaxin with key regions labeled. This conformation retains its compact structure and part of the H3 domain remains in contact with Habc. Rendered in (B) is an overlay of this partially open state with a prior model for the open state of syntaxin, which is characterized by unwinding of the linker region and complete dissociation of H3 from Habc (B). Plotted in (C) are residue-residue distance distributions with values from the open and partially open states indicated by arrows. Both sets of distances agree with the DEER data, suggesting that the syntaxin open state may be more conformationally diverse than previously thought.

Preliminary refinement of the syntaxin conformational ensemble illustrates the importance of heterogeneous ensembles and of having refinement methods that treat them rigorously. Because bias-resampling ensemble refinement is explicitly designed to refine highly heterogeneous conformational ensembles, it significantly out-performs current refinement methods in estimating such ensembles from distributional data. Furthermore, bias-resampling ensemble refinement could be combined with other methods (such as metaynamics,^38^ Rosetta,^39–41^ or other non-MD sampling) to improve their treatment of heterogeneous data. Applied to syntaxin, where the DEER data reveal substantial heterogeneity, our new method uncovers previously unreported heterogeneity in the syntaxin open state. These new conformations may play an important role in mechanisms of SNARE complex assembly.

## ACKNOWLEDGMENTS

The authors thank B. Liang, C. Stroupe, and L. Tamm for many helpful discussions. Computational resources were provided by NCSA Blue Waters, the Advanced Research Computing Services at the University of Virginia, and SNIC and the Center for Parallel Computation at KTH. This work was supported by R01GM115790 to P.M.K., NIH grant PO1 GM072694 to D.S.C, a Wallenberg Academy Fellowship to P.M.K., and a MolSSI Fellowship from the National Science Foundation (ACI-1547580) to J.M.H.

## REFERENCES

(1) Boehr, D. D.; Nussinov, R.; Wright, P. E. The Role of Dynamic Conformational Ensembles in Biomolecular Recognition. Nat. Chem. Biol. 2009, 5 (11), 789–796.

(2) Norman, A. W.; Mizwicki, M. T.; Norman, D. P. G. Steroid-Hormone Rapid Actions, Membrane Receptors and a Conformational Ensemble Model. Nat. Rev. Drug Discov. 2004, 3 (1), 27–41.

(3) Wright, P. E.; Dyson, H. J. Intrinsically Disordered Proteins in Cellular Signalling and Regulation. Nat. Rev. Mol. Cell Biol. 2015, 16 (1), 18–29.

(4) Motlagh, H. N.; Wrabl, J. O.; Li, J.; Hilser, V. J. The Ensemble Nature of Allostery. Nature 2014, 508 (7496), 331–339.

(5) Jimenez, R.; Salazar, G.; Yin, J.; Joo, T.; Romesberg, F. E. Protein Dynamics and the Immunological Evolution of Molecular Recognition. Proc. Natl. Acad. Sci. 2004, 101 (11), 3803–3808.

(6) Mittag, T.; Kay, L. E.; Forman-Kay, J. D. Protein Dynamics and Conformational Disorder in Molecular Recognition. J. Mol. Recognit. 2010, 23 (2), 105–116.

(7) Levin, E. J.; Kondrashov, D. A.; Wesenberg, G. E.; Phillips, G. N. Ensemble Refinement of Protein Crystal Structures: Validation and Application. Structure 2007, 15 (9), 1040–1052.

(8) Bernadó, P.; Mylonas, E.; Petoukhov, M. V.; Blackledge, M.; Svergun, D. I. Structural Characterization of Flexible Proteins Using Small-Angle X-Ray Scattering. J. Am. Chem. Soc. 2007, 129 (17), 5656–5664.

(9) Bonvin, A. M. J. J.; Brünger, A. T. Conformational Variability of Solution Nucelar Magnetic Resonance Structures. J. Mol. Biol. 1995, 250 (1), 80–93.

(10) Jeschke, G. Characterization of Protein Conformational Changes with Sparse Spin-Label Distance Constraints. J. Chem. Theory Comput. 2012, 8 (10), 3854–3863.

(11) van den Bedem, H.; Fraser, J. S. Integrative, Dynamic Structural Biology at Atomic Resolution—It’s about Time. Nat. Methods 2015, 12 (4), 307–318.

(12) Rieping, W. Inferential Structure Determination. Science 2005, 309 (5732), 303–306.

(13) Bonomi, M.; Heller, G. T.; Camilloni, C.; Vendruscolo, M. Principles of Protein Structural Ensemble Determination. Curr. Opin. Struct. Biol. 2017, 42, 106–116.

(14) Bertini, I.; Giachetti, A.; Luchinat, C.; Parigi, G.; Petoukhov, M. V.; Pierattelli, R.; Ravera, E.; Svergun, D. I. Conformational Space of Flexible Biological Macromolecules from Average Data. J. Am. Chem. Soc. 2010, 132 (38), 13553–13558.

(15) Bonomi, M.; Camilloni, C.; Cavalli, A.; Vendruscolo, M. Metainference: A Bayesian Inference Method for Heterogeneous Systems. Sci. Adv. 2016, 2 (1), e1501177.

(16) Bonomi, M.; Camilloni, C.; Vendruscolo, M. Metadynamic Metainference: Enhanced Sampling of the Metainference Ensemble Using Metadynamics. Sci. Rep. 2016, 6 (1), 31232.

(17) Lindorff-Larsen, K.; Best, R. B.; DePristo, M. A.; Dobson, C. M.; Vendruscolo, M. Simultaneous Determination of Protein Structure and Dynamics. Nature 2005, 433 (7022), 128–132.

(18) Chen, J.; Chen, J.; Pinamonti, G.; Clementi, C. Learning Effective Molecular Models from Experimental Observables. J. Chem. Theory Comput. 2018, 14 (7), 3849–3858.

(19) Olsson, S.; Wu, H.; Paul, F.; Clementi, C.; Noé, F. Combining Experimental and Simulation Data of Molecular Processes via Augmented Markov Models. Proc. Natl. Acad. Sci. 2017, 114 (31), 8265–8270.

(20) Olsson, S.; Frellsen, J.; Boomsma, W.; Mardia, K. V.; Hamelryck, T. Inference of Structure Ensembles of Flexible Biomolecules from Sparse, Averaged Data. PLoS ONE 2013, 8 (11), e79439.

(21) Habeck, M.; Rieping, W.; Nilges, M. Weighting of Experimental Evidence in Macromolecular Structure Determination. Proc. Natl. Acad. Sci. 2006, 103 (6), 1756–1761.

(22) Heo, L.; Feig, M. Experimental Accuracy in Protein Structure Refinement via Molecular Dynamics Simulations. Proc. Natl. Acad. Sci. 2018, 115 (52), 13276–13281.

(23) White, A. D.; Voth, G. A. Efficient and Minimal Method to Bias Molecular Simulations with Experimental Data. J. Chem. Theory Comput. 2014, 10 (8), 3023–3030.

(24) Pitera, J. W.; Chodera, J. D. On the Use of Experimental Observations to Bias Simulated Ensembles. J. Chem. Theory Comput. 2012, 8 (10), 3445–3451.

(25) Boomsma, W.; Ferkinghoff-Borg, J.; Lindorff-Larsen, K. Combining Experiments and Simulations Using the Maximum Entropy Principle. PLoS Comput. Biol. 2014, 10 (2), e1003406.

(26) Dawidowski, D.; Cafiso, D. S. Allosteric Control of Syntaxin 1a by Munc18-1: Characterization of the Open and Closed Conformations of Syntaxin. Biophys. J. 2013, 104 (7), 1585–1594.

(27) Marinelli, F.; Faraldo-Gómez, J. D. Ensemble-Biased Metadynamics: A Molecular Simulation Method to Sample Experimental Distributions. Biophys. J. 2015, 108 (12), 2779–2782.

(28) Roux, B.; Islam, S. M. Restrained-Ensemble Molecular Dynamics Simulations Based on Distance Histograms from Double Electron–Electron Resonance Spectroscopy. J. Phys. Chem. B 2013, 117 (17), 4733–4739.

(29) Margittai, M.; Widengren, J.; Schweinberger, E.; Schroder, G. F.; Felekyan, S.; Haustein, E.; Konig, M.; Fasshauer, D.; Grubmuller, H.; Jahn, R.; et al. Single-Molecule Fluorescence Resonance Energy Transfer Reveals a Dynamic Equilibrium between Closed and Open Conformations of Syntaxin 1. Proc. Natl. Acad. Sci. 2003, 100 (26), 15516–15521.

(30) Carr, C. M. The Taming of the SNARE. Nat. Struct. Biol. 2001, 8 (3), 186–188.

(31) Misura, K. M. S.; Scheller, R. H.; Weis, W. I. Three-Dimensional Structure of the Neuronal-Sec1–Syntaxin 1a Complex. Nature 2000, 404 (6776), 355–362.

(32) Jahn, R.; Scheller, R. H. SNAREs — Engines for Membrane Fusion. Nat. Rev. Mol. Cell Biol. 2006, 7 (9), 631–643.

(33) Chen, Y. A.; Scheller, R. H. SNARE-Mediated Membrane Fusion. Nat. Rev. Mol. Cell Biol. 2001, 2 (2), 98–106.

(34) Gerber, S. H.; Rah, J.-C.; Min, S.-W.; Liu, X.; de Wit, H.; Dulubova, I.; Meyer, A. C.; Rizo, J.; Arancillo, M.; Hammer, R. E.; et al. Conformational Switch of Syntaxin-1 Controls Synaptic Vesicle Fusion. Science 2008, 321 (5895), 1507–1510.

(35) Rizo, J.; Südhof, T. C. Snares and Munc18 in Synaptic Vesicle Fusion. Nat. Rev. Neurosci. 2002, 3 (8), 641–653.

(36) Liang, B.; Kiessling, V.; Tamm, L. K. Prefusion Structure of Syntaxin-1A Suggests Pathway for Folding into Neuronal Trans-SNARE Complex Fusion Intermediate. Proc. Natl. Acad. Sci. 2013, 110 (48), 19384–19389.

(37) Wang, S.; Choi, U. B.; Gong, J.; Yang, X.; Li, Y.; Wang, A. L.; Yang, X.; Brunger, A. T.; Ma, C. Conformational Change of Syntaxin Linker Region Induced by Munc13s Initiates SNARE Complex Formation in Synaptic Exocytosis. EMBO J. 2017, 36 (6), 816–829.

(38) Laio, A.; Parrinello, M. Escaping Free-Energy Minima. Proc. Natl. Acad. Sci. 2002, 99 (20), 12562–12566.

(39) Leaver-Fay, A.; Tyka, M.; Lewis, S. M.; Lange, O. F.; Thompson, J.; Jacak, R.; Kaufman, K. W.; Renfrew, P. D.; Smith, C. A.; Sheffler, W.; et al. Rosetta3. In Methods in Enzymology; Elsevier, 2011; Vol. 487, pp 545–574.

(40) Chaudhury, S.; Lyskov, S.; Gray, J. J. PyRosetta: A Script-Based Interface for Implementing Molecular Modeling Algorithms Using Rosetta. Bioinformatics 2010, 26 (5), 689–691.

(41) Hirst, S. J.; Alexander, N.; Mchaourab, H. S.; Meiler, J. RosettaEPR: An Integrated Tool for Protein Structure Determination from Sparse EPR Data. J. Struct. Biol. 2011, 173 (3), 506–514.

