## Supplementary Information and Methods for "Hybrid Refinement of Heterogeneous Conformational Ensembles using Spectroscopic Data"

**Supporting Information for Hybrid refinement of heterogeneous conformational ensembles using spectroscopic data**

### Theory

A more complex formulation of bias-resampling ensemble refinement (BRER) may be used to perform more advanced sampling. Rather than draw each conformation $x$ from the previous conformational estimate $\widehat{\{X\}}_{i-1}$, a history is maintained of $k$ refinement rounds, so that the conformation $x$ is drawn from the union of $\widehat{\{X\}}_{i-1}, \widehat{\{X\}}_{i-2}, \dots, \widehat{\{X\}}_{i-k}$. The conformational estimate $\widehat{\{X\}}_i$ is then obtained as before: the conformations are updated using a biased MD simulation such that the updated estimate $\widehat{\{X\}}_{1\dots i}$ will optimally reproduce $P_{\mathrm{DEER}}(d)$. Just as with the formulation provided in the main text, over the course of multiple rounds of refinement, the conformational estimate $\widehat{\{X\}}$ should yield a distribution $P_{\widehat{\{X\}}}(d)$ that converges on $P_{\mathrm{DEER}}(d)$.

### Methods

#### Molecular dynamics simulations

*Setup and equilibration of syntaxin-1a*

In order to best demonstrate the ability of our method to sample backbone conformational change and rare conformational states, we started all simulations of syntaxin-1a from its closed state. We obtained an initial structure of closed syntaxin by extracting the soluble domain from the crystal structure of syntaxin in complex with Munc-18 (PDB ID 3C98)[1]. The system was solvated with approximately 90,000 TIP3P water molecules and ions were added to obtain a system with 150 mM NaCl and no net charge. Simulations were run in Gromacs[2] using the CHARMM36[3] force field. The system was energy minimized using the steepest-descent integrator for 5000 steps or until the largest force was less than 500 kJ/mol/nm$^2$, whichever came first. A brief 100 ps equilibration was run using NPT conditions using the velocity-rescaling thermostat[4] at 310 K with a 2-ps time constant and pressure maintained at 1 bar using the Parrinello-Rahman barostat[5] with a 10-ps time constant. Covalent bonds were constrained using LINCS, and long-range electrostatics were treated using Particle Mesh Ewald[6]. For each set of ensemble simulations, we generated 50 identical replicas from the equilibrated structure and used these replicas as initial states for production runs.

*Production simulations*

All production simulations were run under the same NPT conditions described above. DEER-derived distance distributions were smoothed with a Gaussian filter. The smoothing parameter σ was chosen to reflect the experimental uncertainty in the fine modes of the DEER-derived distance distributions, 2 Å for all three distributions. Histograms were calculated using 1 Å bins. These distributions were then incorporated into MD simulation using each of three ensemble methods, detailed below. Production simulations were carried out using 50 ensemble members and 5 µs of simulation data were collected for each refinement method except EBMetaD. The reason for this exception is described in "EBMetaD simulations." Simulations were run using Gromacs[2] and the *gmxapi* Python API[7], which permits introduction of user-defined biasing potentials.

*BRER simulations*

To sample the syntaxin conformational ensemble, we performed five iterations of BRER for each of 50 ensemble members. Each iteration is performed as follows: first, one target distance is chosen from each of the smoothed DEER distributions, then a linear biasing potential

$$U_{\text{bias}} = \sum_{n=1}^{N_{\text{distributions}}} \alpha_i \frac{d_{\text{MD}}^{(n)}}{d_{\text{target}}^{(n)}}$$

is applied to drive the simulation distance to the target.

Convergence to the target is achieved in two phases. During the training phase, the Hamiltonian coupling constants {α} are learned for each target using a modified version of the method described by White and Voth.[8] Each constant α is updated every 50 ps according to

$$\alpha_\tau = \alpha_{\tau-1} - \eta_\tau g_\tau,$$

where $\eta$ is the learning rate and $g$ is the gradient:

$$g_\tau = -2\beta\left(\frac{\langle d_{\text{MD}}\rangle_\tau}{d_{\text{target}}} - 1\right)\left(\langle d_{\text{MD}}^2\rangle_\tau - \langle d_{\text{MD}}\rangle_\tau^2\right),$$

$$\eta_\tau = \frac{A}{\sqrt{\sum_{i=1}^{\tau} g_i}}$$

At the end of the training phase, we select the maximum value of α to prevent underestimating α if the $i^{\text{th}}$ degree of freedom converges much faster than the others. During the convergence phase, the simulation is restarted from the beginning of the iteration and a time independent potential (α fixed) is applied until the simulation converges to the target. The parameter *A* was chosen so as to achieve convergence between 1-5 ns (*A*=150β). Once the simulation has converged to the target, a 20 ns production run is performed to relax the remaining degrees of freedom. The full procedure is then repeated, beginning with random resampling from the DEER distributions.

A python package to run BRER ensemble simulations is available at https://github.com/jmhays/run_brer and documentation can be found at https://run-brer.readthedocs.io/en/latest/. A singularity container is also available at https://singularity-hub.org/collections/1761.

*EBMetaD simulations*

EBMetaD simulations were implemented using the same modified version of Gromacs and gmxapi[7] version as the BRER simulations. Because the EBMetaD potential is ill-defined in regions of zero probability, we add a small uniform prior to all experimental distributions: using the same notation as Marinelli and Faraldo-Gomez,[9] the modified EBMetaD potential is

$$V(\xi, t) = \sum_{t'=\tau,2\tau\ldots}^{t} \frac{w\exp\{-[\xi - \xi^f(X_{t'})]^2/2\sigma^2\}}{\exp\{S_\rho\}\left(\rho_{\exp}[\xi^f(X_{t'})] + \delta_{\text{uniform}}\right)}$$

where we have added the term $\delta_{\text{uniform}}$. As $\delta_{\text{uniform}}$ increased, the simulations become more numerically stable, but the solution approaches standard metadynamics. Therefore, $\delta_{\text{uniform}}$ should be chosen carefully so as to maintain information about the DEER distributions but still produce stable simulations. We selected $\delta_{\text{uniform}}$=0.1. Even with this choice of $\delta_{\text{uniform}}$, the method exhibited a high rate of stochastic failure: all 50 ensemble members failed in the range of 10ns-150 ns of simulation time. Because of this, we were only able to collect ~ 3μs of data.

This version of the EBMetaD method is available at https://github.com/jmhays/run_ebmetad. A singularity container is also available at https://www.singularity-hub.org/collections/1994.

*Restrained-ensemble simulations*

Restrained-ensemble biasing potentials previously developed by Roux[10-11] were applied to match MD distance histograms to DEER-derived distance distributions. Refinement was performed via restrained-ensemble simulation using a modified version of Gromacs 5.2 available at https://github.com/kassonlab/restrained-ensemble. This method exhibits numerical instabilities when distributions are very tightly peaked, such as when an ensemble is started from copies of a single initial state. Thus, we initially used a very broad smoothing parameter, σ=10 Å, for both the MD and DEER-derived distributions, which we modified to σ=1 Å once the ensemble had sampled enough of the distribution to be stable for small σ. Distance data were collected for all ensemble members for a period of 100 ps followed by an update of the biasing potential with a spring constant of K=100 kJ/mol/$nm^2$. Additionally, a boxcar averaging filter was applied so that the simulation distance distributions were calculated using the last 10 ns of data. These modifications were implemented in order to obtain sufficient sampling for generating the MD distance distributions as previously described in Hays et al.[12]

### Calculation of final distributions and Jensen-Shannon divergence

For each ensemble, production simulations were sampled at 500 ps intervals and distances between the Cβ of each residue-residue pair measured by DEER were calculated using MDAnalysis[13]. The distributions plotted in Fig. 3 of the main text were calculated using a Gaussian filter with smoothing parameter σ = 2 Å and 1 Å bins for all three distributions. This was done for consistency with the experimental data, which was also smoothed with σ = 2 Å and 1 Å bin width. Jensen-Shannon divergence was calculated using the smoothed experimental and simulation distributions.

### Conformational ensemble analysis

We clustered the BRER-refined structures as follows: final, relaxed structures from each stochastic-resampling iteration were collected and the distances between the Cβ of each residue-residue pair measured by DEER were calculated using MDAnalysis[13]. Distances were calculated from Cα for glycine residues. We performed k-means clustering on these distance coordinates for a broad range of cluster numbers (2 – 50 clusters). We selected the smallest cluster number (20) for which the average intra-cluster RMSD was substantially higher than the average inter-cluster RMSD (7 Å and 9 Å, respectively). Clusters were classified as “open” if the 52/210 distance was > 40 Å. The structure rendered in Fig 4 of the main text is the centroid of the most populated open cluster.

### References


1. Burkhardt, P.; Hattendorf, D. A.; Weis, W. I.; Fasshauer, D., Munc18a controls SNARE assembly through its interaction with the syntaxin N-peptide. *The EMBO journal* **2008,** *27* (7), 923-33.
2. Pronk, S.; Pall, S.; Schulz, R.; Larsson, P.; Bjelkmar, P.; Apostolov, R.; Shirts, M. R.; Smith, J. C.; Kasson, P. M.; van der Spoel, D.; Hess, B.; Lindahl, E., GROMACS 4.5: a high-throughput and highly parallel open source molecular simulation toolkit. *Bioinformatics* **2013,** *29* (7), 845-54.
3. Huang, J.; MacKerell, A. D., Jr., CHARMM36 all-atom additive protein force field: validation based on comparison to NMR data. *J Comput Chem* **2013,** *34* (25), 2135-45.
4. Bussi, G.; Donadio, D.; Parrinello, M., Canonical sampling through velocity rescaling. *J Chem Phys* **2007,** *126* (1), 014101.
5. Parrinello, M.; Rahman, A., Strain Fluctuations and Elastic-Constants. *Journal of Chemical Physics* **1982,** *76* (5), 2662-2666.

6. Darden, T.; York, D.; Pedersen, L., Particle Mesh Ewald - an N.Log(N) Method for Ewald Sums in Large Systems. *Journal of Chemical Physics* **1993,** *98* (12), 10089-10092.
7. Irrgang, M. E.; Hays, J. M.; Kasson, P. M., gmxapi: a high-level interface for advanced control and extension of molecular dynamics simulations. *Bioinformatics* **2018,** *34* (22), 3945-3947.
8. White, A. D.; Voth, G. A., Efficient and Minimal Method to Bias Molecular Simulations with Experimental Data. *J Chem Theory Comput* **2014,** *10* (8), 3023-30.
9. Marinelli, F.; Faraldo-Gomez, J. D., Ensemble-Biased Metadynamics: A Molecular Simulation Method to Sample Experimental Distributions. *Biophys J* **2015,** *108* (12), 2779-82.
10. Roux, B.; Islam, S. M., Restrained-ensemble molecular dynamics simulations based on distance histograms from double electron-electron resonance spectroscopy. *J Phys Chem B* **2013,** *117* (17), 4733-9.
11. Islam, S. M.; Stein, R. A.; McHaourab, H. S.; Roux, B., Structural refinement from restrained-ensemble simulations based on EPR/DEER data: application to T4 lysozyme. *J Phys Chem B* **2013,** *117* (17), 4740-54.
12. Hays, J. M.; Kieber, M. K.; Li, J. Z.; Han, J. I.; Moremen, K. W.; Columbus, L.; Kasson, P. M., Refinement of highly flexible protein structures using simulation-guided spectroscopy. *Angew Chem Int Ed Engl.* **2018,** *57* (52), 17110-17114.
13. Michaud-Agrawal, N.; Denning, E. J.; Woolf, T. B.; Beckstein, O., MDAnalysis: A toolkit for the analysis of molecular dynamics simulations. *J. Comput. Chem.* **2011,** *32* (10), 2319--2327.
